# Identification, expression analysis and molecular modeling of *Iron deficiency specific clone 3* (*Ids3*) like gene in hexaploid wheat

**DOI:** 10.1101/244392

**Authors:** Priyanka Mathpal, Upendra Kumar, Anuj Kumar, Sanjay Kumar, Sachin Malik, Naveen Kumar, H. S. Dhaliwal, Sundip Kumar

## Abstract

Graminaceous plants secrete iron (Fe) chelators called mugineic acid family phytosiderophores (MAs) from their roots for solubilisation and mobilization of unavailable ferric (Fe^3+^) ions from the soil. The hydroxylated forms of these phytosiderophores have been found more efficient in chelation and subsequent uptake of minerals from soil which are available in very small quantities. The genes responsible for hydroxylation of phytosiderophores have been recognized as iron deficiency-specific clone 2 (*Ids2*) and iron deficiency-specific clone 3 (*Ids3*) in barley but their presence is not reported earlier in hexaploid wheat. Hence, the present investigation was done with the aim:(i) to search for the putative *Hordeum vulgare Ids3* (*HvIds3*) ortholog in hexaploid wheat, (ii) physical mapping of *HvIds3* ortholog on wheat chromosome using cytogenetic stocks developed in the background of wheat cultivar Chinese Spring and (iii) to analyze the effect of iron starvation on the expression pattern of this ortholog at transcription level. In the present investigation, a putative ortholog of *HvIds3* gene was identified in hexaploid wheat using different bioinformatics tools. Further, protein structure of TaIDS3 was modelled using homology modeling and also evaluated modelled structure behavior on nanoseconds using molecular dynamics based approach. Additionally, the ProFunc results also predict the functional similarity between the proteins of *HvIds3* and its wheat ortholog (*TaIds3*). The physical mapping study with the use of cytogenetic stocks confines *TaIds3* in the telomeric region of chromosome 7AS which supports the results obtained by bioinformatics analysis. The relative expression analysis of *TaIds3* indicated that the detectable expression of *TaIds3* induces after 5^th^ day of Fe-starvation and increases gradually up to 15^th^ day and thereafter decreases till 35^th^ day of Fe-starvation. This reflects that Fe deficiency directly regulates the induction of *TaIds3* in the roots of hexaploid wheat.

## Introduction

Iron (Fe) deficiency, a major abiotic stress, especially in calcareous soils (~30% of the world’s cultivated soils) with extremely low solubility of Fe (owing to high pH levels) reduces crop yield. The low availability of Fe results into its poor uptake by plants, which causes severe yield losses. Further, the poor uptake of other minerals by plants also reduces the overall mineral content in grains, which are essential to meet the required human dietary needs (Cakmak 2008).

The manifestation of Fe deficiency in calcareous soils has been recognised as Fe-chlorosis and lime-induced chlorosis. Under adverse conditions of Fe deficiency, graminaceous plants secrete Fe-chelators called mugineic acid family phytosiderophores from their roots (Takagi 1978). Mugineic acid family phytosiderophores (MAs) are highly effective in solubilisation and mobilization of Fe in calcareous soil (Treeby et al. 1989) and are involved in the uptake of Fe through roots (Romeheld and Marschner 1990; vonWiren et al. 1995) to acquire sparingly soluble Fe as Fe^3+^-MAs complex through the use of Fe^3+^-MA transporters. The secretion of MAs increase under deficiency of Fe and is correlated with plant’s tolerance to Fe deficiency. The quantity and different forms of phytosiderophores secreted by roots play an important role in providing tolerance to plants in Fe-deficient condition. The hydroxylated forms of phytosiderophores like 3-hydroxymugineic acid (HMA) and 3-epihydroxymugineic acid (epiHMA) have been found more efficient in chelation and subsequent uptake of trace minerals (vonWiren et al. 2000). In *Hordeum vulgare*, the genes responsible for hydroxylation of phytosiderophores have been recognized as *iron deficiency-specific clone 2* (*Ids2*) and *iron deficiency-specific clone 3* (*Ids3*) (Nakanishi et al. 2000). The *Ids3* gene encodes an enzyme dioxygenase which hydroxylates the C-2ʹ position of 2ʹ-deoxymugineic acid (DMA) and 3-epihydroxy-2ʹ-deoxymugineic acid (epiHDMA) and converts DMA to MA or epiHDMA to epiHMA, while *Ids2* hydroxylates C-3ʹ position of MA and DMA and converts MA to epiHMA or DMA to epiHDMA. MA, epiHDMA and epiHMA are more stable under mildly acidic conditions and might be more favourable for inter mineral translocation (Kobayashi and Nishizawa 2012). As reported in publicly available sequence databases, *Ids2* is having 3,411 bp long coding sequence which encodes a protein of 338 amino acids (http://www.uniprot.org/uniprot/Q40061) while *Ids3* have 4,904 bp long coding sequence which encodes a protein of 339 amino acids (http://www.uniprot.org/uniprot/Q40062). The expression of *Ids3* is always much stronger than that of *Ids2* in Fe-deficient condition, indicating that initially DMA changes into MA predominantly (Nakanishi et al. 2000).

The biosynthetic pathway for mugineic acid production is well documented in graminaceous crops (Ma and Nomoto 1993). All MAs share the pathway from methionine to DMA and the DMA is then converted to other MAs like HMA, epiHMA and epiHDMA. The pathway up to DMA synthesis is conserved in all graminaceous plants and the variation is in the type of phytosiderophores synthesized in the further hydroxylation steps (Takahashi 2003). It is reported that wheat cultivar Chinese Spring used in the present study produces only one phytosiderophore, namely DMA (non-hydroxylated form) whereas barley cv. Betzes and rye each synthesizes four types of MAs i.e. DMA, MA, epi-HDMA and epi-HMA (Nakanishi et al. 2000; Mori et al. 1990; Neelam et al. 2011). It has been reported that wheat and its wild progenitors do not have the ability to secrete MA and HMA from the beginning of their evolution because they lack the ability to secrete MA and HMA as a result of the absence of IDS3 enzyme (Singh et al. 1973). Therefore, no efforts were made in the past to study *Ids3* gene in wheat genome.

In view of the above, the present investigation was aimed at the identification and physical mapping of the wheat ortholog of *Ids3* gene and to analyze its expression pattern under Fe-deficient conditions in hexaploid wheat.

## Materials and Methods

### Plant Growth Conditions

Seeds of wheat cultivar Chinese Spring were surface sterilized and soaked on filter paper with distilled water and incubated in dark at 25 °C until germination. Then the three-days old seedlings were transplanted and cultured hydroponically in 2.0 litre plastic boxes with continuous air flow under Fe-sufficient (+Fe) and Fe-deficient (−Fe) conditions. The 1X concentration of the nutrient solution for hydroponic culture consisted of 2.0mM Ca(NO_3_)_2_, 0. 7mM K_2_SO_4_, 0.1mM KCl, 0.1mM KH_2_PO_4_, 0.50mM MgSO_4_, 0.15mM Fe^3+^-EDTA, 10µM H_3_BO_3_, 0.5μΜ MnSO_4_, 0.5μΜ ZnSO_4_, 0.2μΜ CuSO_4_ and 0.01μΜ (ΝΗ_4_)_6_Μo_7_O_24_. The nutrient solution was replaced once per week and its pH adjusted to 5.5 to 5.6 daily with 1N HCL. For expression study under Fe starvation, wheat cultivar Chinese Spring was grown at 23–25 °C under a 16/8 h light (~160 μmol/m^2^.s)/dark regimen.

### Identification of *Hordeum vulgare Ids3* homolog through *in-silico* approach

*In-silico* analysis was performed to identify the putative ortholog(s) of *HvIds3* gene in hexaploid wheat. Full length genomic sequence of *HvIds3* gene (Accession Number AB024058.1) was retrieved from NCBI (National Center of the Biotechnology Information) (http://www.ncbi.nlm.nih.gov/) in FASTA format. Similarity searches were carried out using genomic sequence of *HvIds3* as a query against the wheat chromosome survey sequences available at http://urgi.versailles.inra.fr/using the BLAST (Basic Local Alignment Search Tool) search engine (http://www.ncbi.nlm.gov/BLAST). The matching wheat contigs having good coverage (>85%), identity (>70%) and e-value (<e-10) were downloaded. The exon/intron structure prediction in the selected wheat contigs was carried out using FGENESH+ (http://linux1.softberry.com/berry.phtml) program. Conserved domain search (CD-Search) program was used to predict functional domain(s) in the sequences of *HvIds3* and its wheat ortholog *TaIds3* (http://www.ncbi.nlm.nih.gov/Structure/cdd/cdd.shtml).

Multiple sequence alignment (MSA) between *HvIds3* and three copies of *TaIds3* (*TaIds3_7AS, TaIds3_7BS*, and *TaIds3_7DS*) was performed by using CLUSTAL W2 (www.ebi.ac.uk/Tools/msa/clustalw2) available on EMBL-EBI (www.ebi.ac.uk). The 3D structure of proteins (*HvIds3* and *TaIds3*) was predicted using homology modeling based Swiss-Model server (http://swissmodel.expasy.org/). Flexible structure AlignmenT by Chaining AFPs (Aligned Fragment Pairs) with Twists (FATCAT) structure alignment tool (http://fatcat.burnham.org/) was used for the alignment of 3D structure model of proteins. ProFunc (http://www.ebi.ac.uk/thornton-srv/databases/ProFunc/) was used to predict the biochemical function of *TaIds3* based on the predicted 3D model. Sub cellular localization of *HvIds3* and *TaIds3* was predicted with TargetP 1.1 (http://www.cbs.dtu.dk/services/TargetP/) and PLANT-mPLoc (http://www.csbio.sjtu.edu.cn/cgi-bin/PlantmPLoc.cgi), respectively.

### Cytogenetic Stocks

To facilitate the mapping of wheat ortholog of *Ids3* gene to the individual chromosome, chromosome arm and sub chromosomal location (i.e. deletion bin), the following cytogeneticstocks in the background of wheat cultivar Chinese Spring were used: (i) 22 nullisomic-tetrasomic (NT) lines (Sears 1954; Sears 1964) (ii) two ditelosomic (Dt) lines i.e. 7AS and 7AL (Sears and Sears 1978) and (iii) 8 homozygous deletion (Del) lines of chromosome 7AS (Endo and Gill 1996). Seeds of NT, Dt and Del lines were procured from the Wheat Genetic and Genomic Resources Center at the Kansas State University, Manhattan, USA and multiplied in the greenhouse facility at the G. B. Pant University of Agriculture & Technology at Pantnagar, India.

### DNA isolation and primer designing

The genomic DNA from fresh leaves of cytogenetic stocks and wheat cultivar Chinese Spring was extracted using CTAB (Cetyltrimethyl ammonium bromide) method with some modifications (Maroof et al. 1994). To confirm the chromosomal location of *TaIds3*, a pair of 7AS chromosome specific primer was designed using primer blast software (http://www.ncbi.nlm.nih.gov/tools/primer-blast/) (Table 1).

**Table 1.**
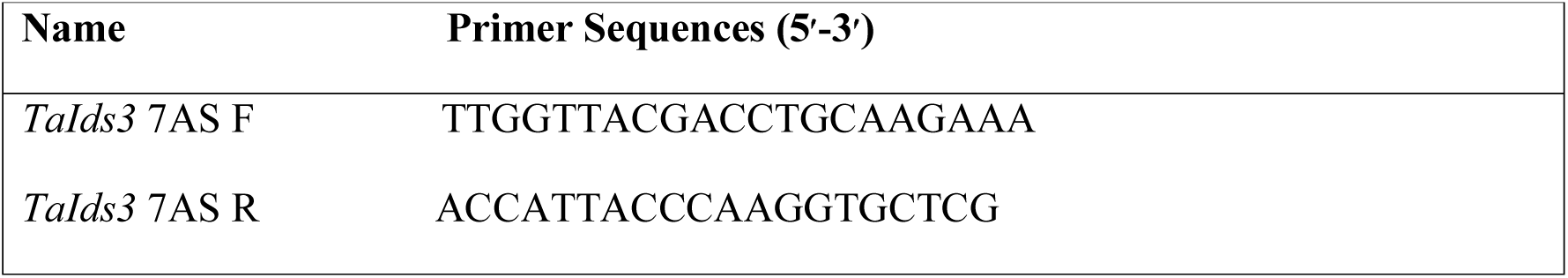
Primers used for chromosomal location of *TaIds3*.

### Polymerase Chain Reaction

Polymerase chain reaction (PCR) amplification was carried out in a volume of 20 µl in Applied Biosystems (Veriti 96 well) thermal cycler. The reaction mixture contained 80–100 ng of template DNA, 1X PCR buffer, 1.5mM MgCl_2_, 0.25mMdNTP, 500nM of each PCR primer (forward and reverse) and 1.0 U *Taq* DNA polymerase (New England Biolabs). PCR cycling conditions were as follows: 95 °C for 2.30 min followed by 35 cycles of 95 °C for 50 s, 56 °C for 40 s and 68 °C for 1.0 min with a final extension of 68 °C for 5 min. The PCR products were resolved using 10% polyacrylamide gel electrophoresis (PAGE) and visualized with silver staining (Tegelstrom 1992).

### RNA extraction and real time reverse-transcription PCR (RT-PCR) analysis

100 mg root samples were harvested at 5, 10, 15, 25 and 35 days after the plants were exposed to +Fe and −Fe conditions and used immediately for RNA isolation. Root tissues were frozen in 2 ml micro centrifuge tubes and disrupted under frozen conditions using two stainless steel beads (5 mm diameter) in the Tissuelyser (Qiagen, USA). Total RNA was extracted from the disrupted tissue using the GeneJET Plant RNA purification kit (Fermentas Life Sciences) according to the manufacturer’s protocol in duplicates. The quantity and quality of the total RNA were evaluated using a nanophotometer (Nanodrop 1000, Thermo-scientific) and visualized by 1% agarose gel electrophoresis. Approximately 1 µg of total RNA was reverse transcribed to cDNA in 20 µL reaction using oligo-dT and M-MLV reverse transcriptase (Fermentas Life Sciences). Real-time PCR primer was designed to amplify a 100–200 bp fragment in untranslated regions. The primer was designed from the conserved region of mRNA of all predicted TaIds3 orthologs using Primer Express® Software version 3.0 (Applied Biosystems) (Table 2).

**Table 2.**
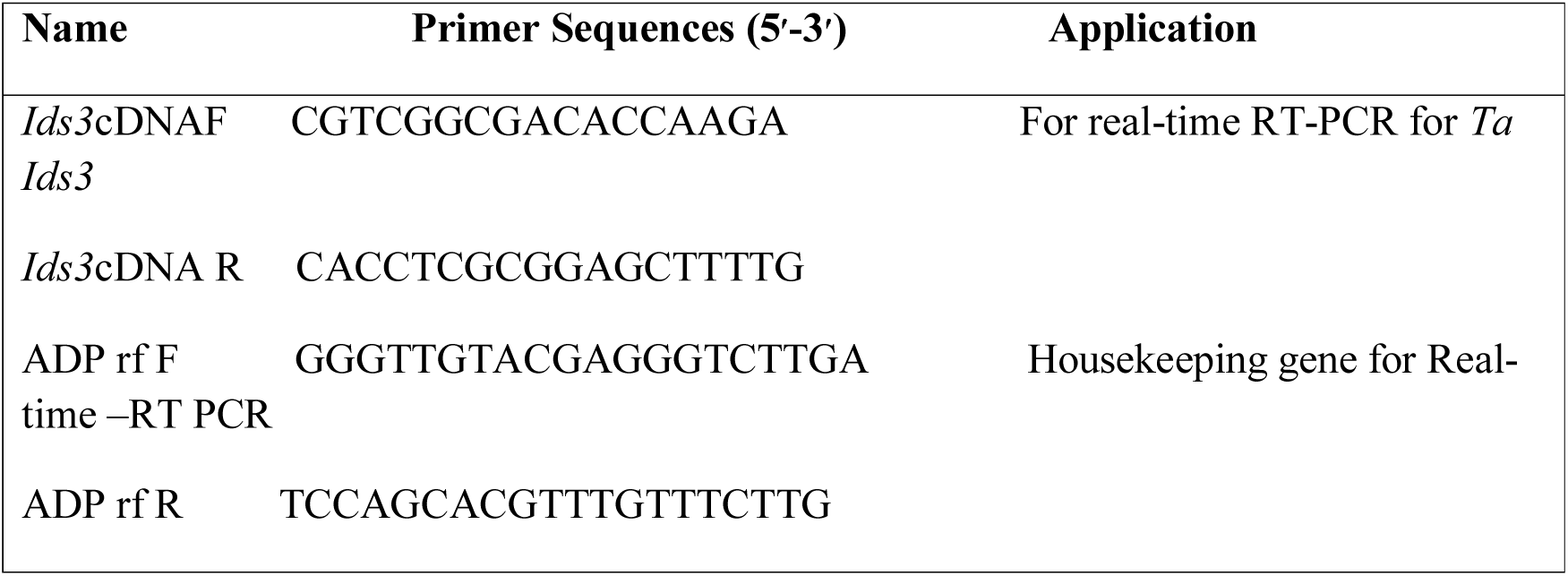
Primers for relative expression analysis of *TaIds3*.

Real-time RT-PCR was performed with a Step One Plus Real-Time PCR System using SYBR Select Master Mix (Applied Biosystems, USA). Reactions were performed in a total volume of 10 µL with 0.2µM of each primer, 1× SYBR premix, 50ng cDNA and ddH_2_O. Reactions were cycled under the following conditions: initial denaturation at 95 °C for 2 min, followed by 40 cycles composed of 15 seconds denaturation at 95 °C and 1.0 min of annealing/extension at 60 °C. To verify specific amplification, melting curve analysis was performed at 65 °C to 95 °C with the fluorescence continuously being monitored. Data were analysed via 2^−ΔΔC^_T_ method with the v2.3 Software (Applied Biosystems, USA) (Livak and Schmittgen, 2001) and the expression level of housekeeping gene *TaADP* ribosylation factor was used as an internal control (Poalacci et al. 2009) (Table 1). For all real-time PCR analysis, two biological replicates were used and three technical replicates were performed for each biological replicate.

### Molecular Dynamics simulations

Molecular dynamics (MD) simulations have become one of the most important techniques in biophysics and helpful to understanding the behaviour of biological macromolecules on nanosecond to microsecond time scales (Gajula et al. 2017). To check the stability of modelled structure of IDS3, we performed molecular dynamics (MD) simulation carried out by Desmond v3.6 (Shiva Kumar et al. 2010; Guo Z et al. 2010; Kevin et al. 2006) implemented in Schrodinger-maestro2017 (Schrödinger Release 2017–2). Protein model preparation done by assigning bond orders, hydrogen atoms added in protein model with neutral pH 7.0 applied by PROPKA (Hui et al. 2005) method. Total 5307 TIP3P (Jorgensen et al. 1983) water solvent atoms were added in truncated octahedron simulation box under periodic boundary conditions and shape and size was set at 10.0Å buffered distance. To neutralize the simulation system electrically, added Na+/Cl- ions to stabilize box charge and were place to randomly in the solvated system. After preparing simulation box (Box size=19882) was proceed to energy minimization by applied 800 steps of steepest descent algorithm followed 2000 steps of conjugate gradient algorithm with 120.0 kcal/mol/Å convergence threshold energy under NPT ensemble. The 300K temperature was applied using Nose-Hoover (Nosé, 1984; Hoover and William, 1984) chain method and 1 atmospheric pressure applied by Martyna-Tobias-Klein (MTK) (Martyna et al. 1994) Barostat method. For both ensemble classes (NPT and NVT) OPLS2005 (Jorgensen et al. 1996; Kaminski et al. 2001) force field were used. Finally, simulation system was set in relaxation state and run for 20nsec.Trajectory recorded in each 4.8psec.

### Simulation Trajectory analysis

The 20 nsec MD simulation trajectory was analyses by simulation event and interaction diagram program module available in Desmond v3.6. For protein stability analysis we select root-mean-square deviation (RMSD) value and root mean square fluctuation (RMSF) selected for determination of fluctuation/thermal motion in protein residues during simulation. We analyze the protein contribution of secondary structure elements (SSE) during simulation in structure stabilization.

## Results

### *In-silico* mapping and functional annotation

Putative wheat ortholog of *HvIds3* gene was identified using different bioinformatics based algorithms. The results suggested the presence of 2OG-FeII Oxy domain (Pfam Id-CL0029) belonging to DIOX protein family (Fig. S1) in wheat ortholog of *HvIds3*. Further, *HvIds3* sequence was aligned against wheat survey sequences available on IWGSC database. BLASTN results showed 74% sequence similarity with IWGSC_chr7AS_ab_k71 contig of chromosome 7AS, 88% sequence similarity with IWGSC_chr7BS_ab_k71 contig of chromosome 7BS and 78% with IWGSC_chr7DS_ab_k71 contig of chromosome 7DS in wheat genome. The predicted structure of 7AS and 7BS contained 4 exons and the structure belonging to 7DS contained 3 exons. *TaIds3_7AS* and *TaIds3_7BS* coded for 339 and 309 amino acid long protein respectively, while *TaIds3_7DS* coded 242 amino acids length protein which was comparable to the protein of the known gene *HvIds3* of barley. Further MSA results predicted by CLUSTALW2 revealed that *TaIds3_7AS* shared high similarity with *HvIds3* as compared to *TaIds3_7BS* and *TaIds3_7DS* (Fig. S2).With the aim to confirm the chromosomal location of *TaIds3* in hexaploid wheat genome, when contig IWGSC_chr7AS_ab_k71 was used to BLAST against *Triticum aestivum* DNA database in Ensembl plants database, it gave a hit on telomeric region of 7AS.

Out of three predicted wheat orthologs, only *TaIds3* on 7AS had complete gene structure (Fig. 1A). Transcription start site (TSS) for *TaIds3* on 7BS and 7DS could not be predicted. CD search for these predicted genes showed that *TaIds3_7AS* with 2OG-FeII Oxy domain belonging to DIOX protein family similar to the gene *HvIds3*, while this domain was not fully present in *TaIds3_7BS* and *TaIds3_7DS* (Fig. 1B).

**Fig. 1.**
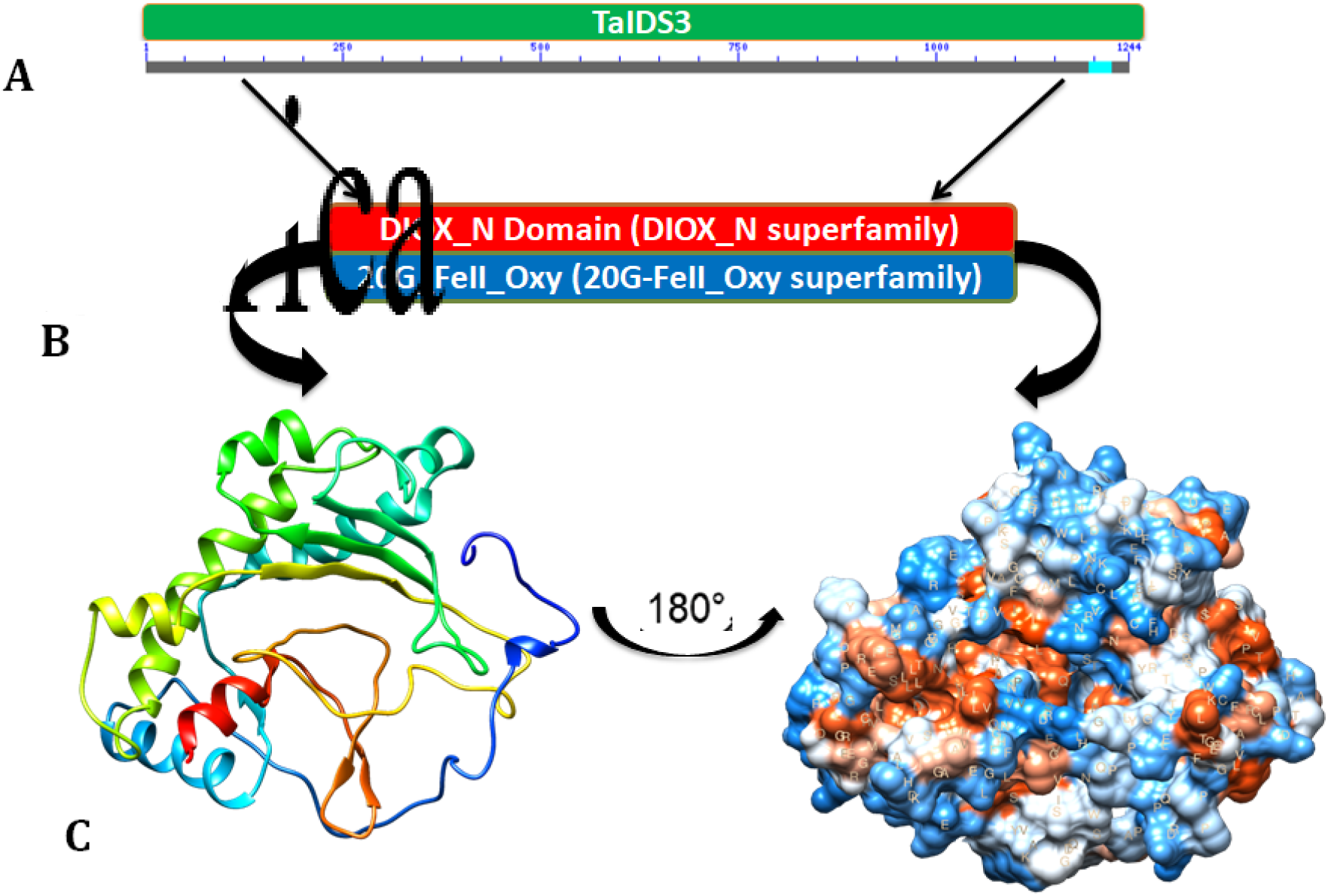
**A)** TaIDS gene structure; (**B**) representation of functional domains and their respective superfamiles; (**C**) interactive ribbon view and surface representation of TaIDS3 protein: Right: view from back of molecule via vertical rotation by 180°. Figure was prepared using UCSF-Chimera.

### Protein Structure Modeling and Phylogeny Analysis

Swiss-Model software (homology based algorithm) was used to generate 3D model wheat proteins encoded by *Ids3* gene based on 1w9yA template from the Protein Data Bank (PDB) at Root Mean Square Deviation (RMSD) 2.0. UCSF-CHIMERA visualized the different chemical shapes of modelled *TaIDS3* (Fig. 1C). The alignment of the structure of the proteins encoded by *HvIds3* and *TaIds3_7AS* using FATCAT superposition showed 57.25% similarity between the *HvIds3* and *TaIds3* proteins (Fig. S3). ProFunc server was used to determine the biochemical function of *Ids3* in wheat showing catalytic activity (59.07) and oxidoreductase activity (49.52) (Table 1). To identify the sub cellular localization of *TaIds3* using machine-learning program PLANT-mPLoc suggests the cytoplasmic localization of *HvIds3, TaIds3_7AS, TaIds3_7BS*, and *TaIds3_7DS*, (Table 2).

The phylogenetic tree generated the individual branches for *TaIds3, HvIds3, HvIds2*, and *OsIds2* (Fig. 2). *TaIds3* gene clusters shared the close evolutionary relationship with *HvIds3*, while *OsIds2* and *OsIds3* shared the relationship with *HvIds2*.

**Fig 2.**
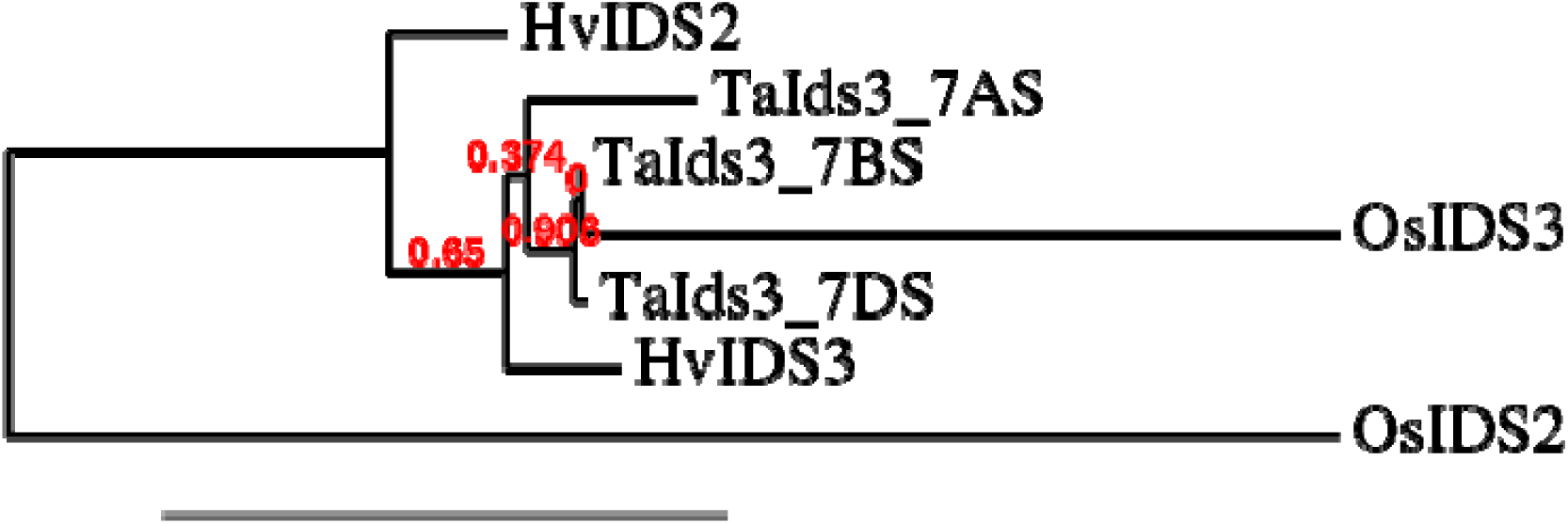
Neighbour-joining phylogenetic tree of the *Ids3* and *Ids2* members. The tree was constructed with the amino acid sequences of *Ids3* and *Ids2* proteins from Barley (Hv), Rice (Os) and Wheat (Ta) using the neighbour-joining method in MEGA 6.0 software. A bootstrap with 1000 repetitions was included.

### Physical mapping of *TaIds3* on 7AS chromosome

In order to confirm the chromosomal location of *TaIds3* in hexaploid wheat genome, the genomic DNA of the cytogenetic stocks developed in Chinese Spring wheat background, was amplified with a pair of 7AS specific primers of *TaIds3* gene. The PCR products of 22 NT lines and di-telocentric lines of 7A are illustrated in Figure 3. The PAGE analysis of the PCR products resolved that a single fragment of *TaIds3* was amplified in 20NT lines, while there was no amplification in N7A-T7B and N7A-T7D. The amplification of the fragment in all 20 NT lines and absence in N7A-T7B and N7A-T7D indicates the presence of this fragment on chromosome 7A (Fig. 4). Further, to localize the fragment on the specific arm of 7A chromosome, the genomic DNA of Dt7AS and Dt7AL was amplified with the same primers. The PAGE analysis of the PCR product resolved that there was no amplification in Dt7AL while Dt7AS line gave the amplification with 7AS specific primers (Fig. 3). These results indicate the location of the gene on short arm of 7A chromosome. Further, the genomic DNA of deletion lines of 7AS didn’t give any amplification. Therefore, it was confirmed that *TaIds3* is located in the telomeric region of 7AS. On the basis of the above results, the location of *TaIds3* gene is illustrated in Fig. 4.

**Fig. 3.**
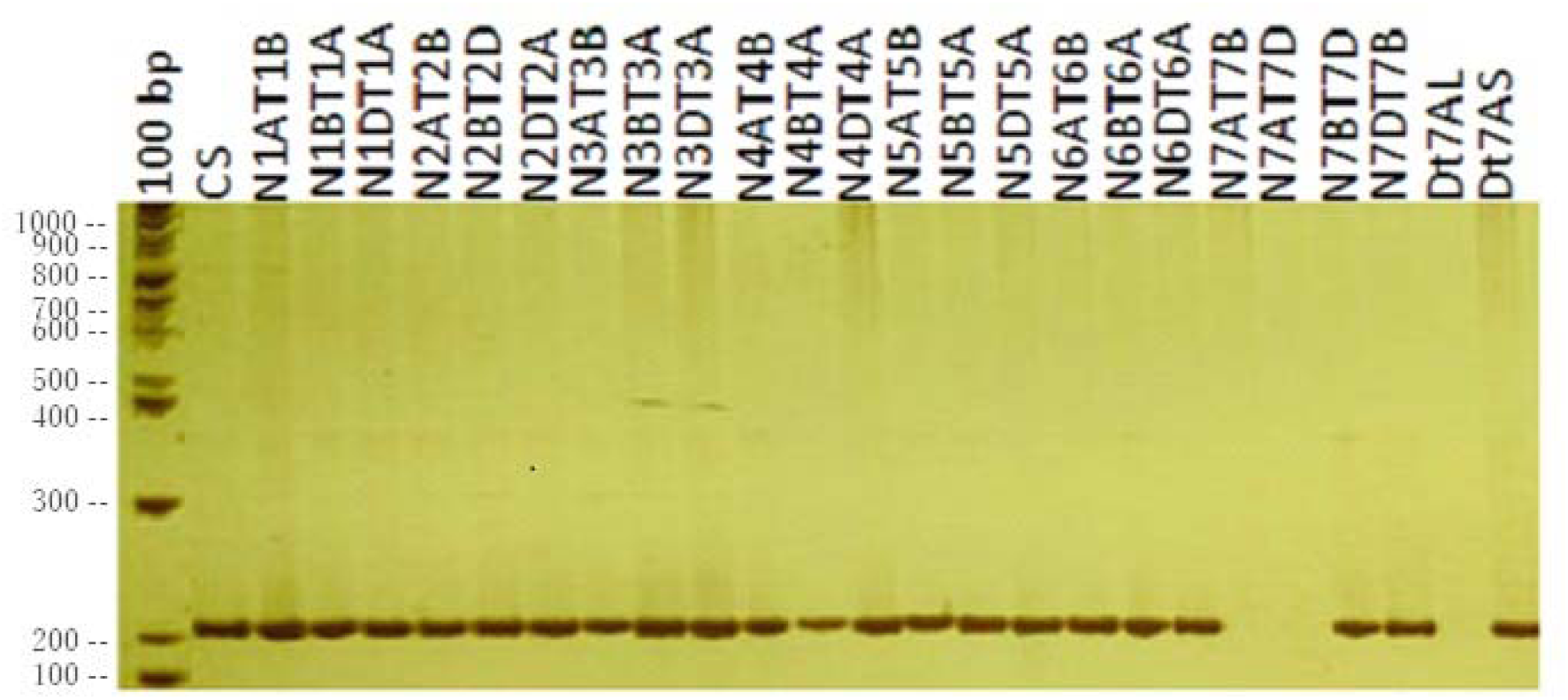
Polyacrylamide gel electrophoresis analysis of the PCR products of *TaIds3* amplified using nullisomic-tetrasomic and ditelosomic lines of *T. aestivum* cv. Chinese Spring.

**Fig. 4.**
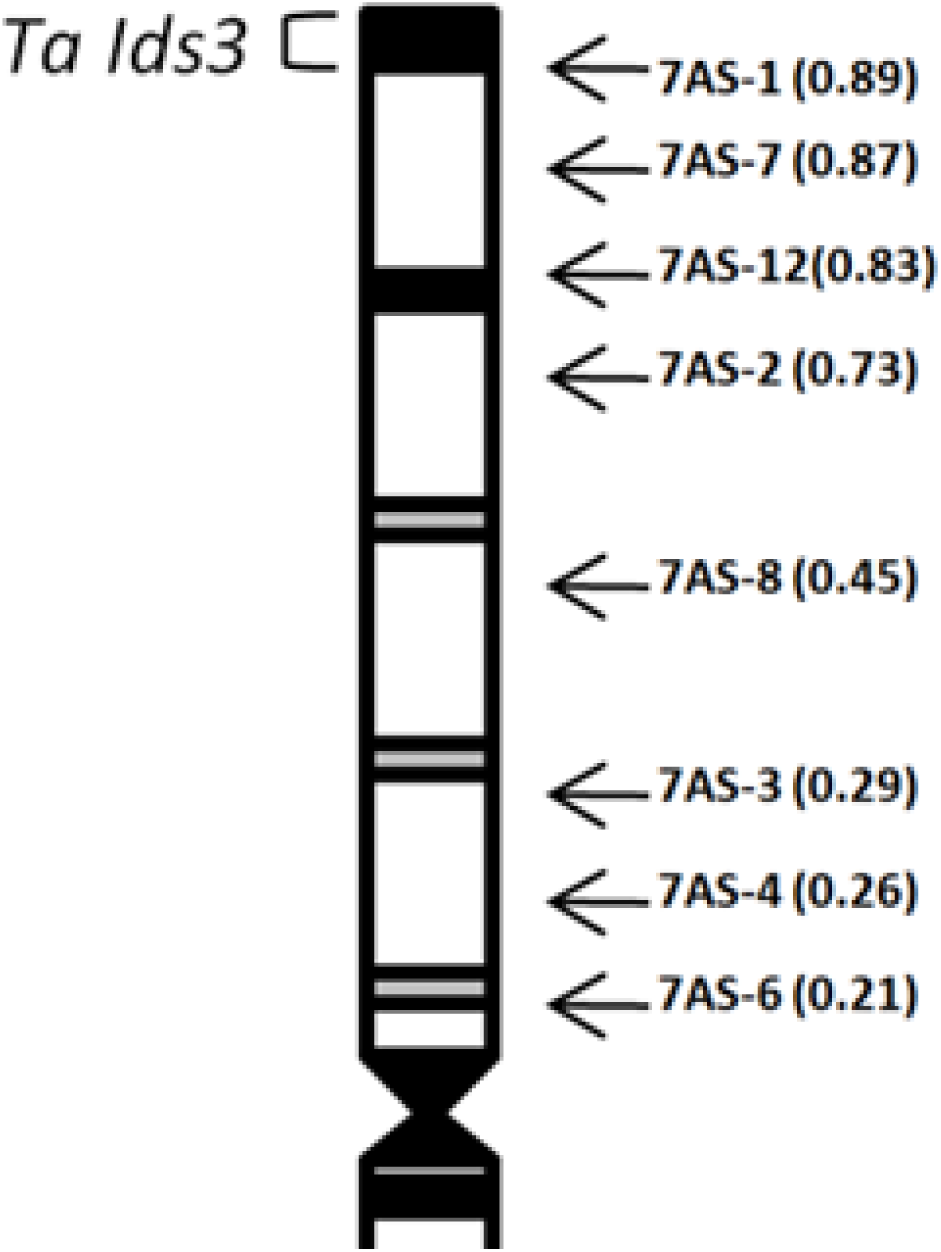
Physical map of Chinese Spring chromosome 7A. The identification of deletion line fraction arm length (FL) values breakpoints is indicated on the right in parentheses. The telomeric bin of 7AS shows the putative location of the *TaIds3* gene.

### Expression analysis under Fe-starvation

The results of Real-time RT-PCR showed that the detectable expression of *TaIds3* was started on 5^th^day (2.5 fold) after Fe-starvation. In response to Fe-deficiency, the relative expression was increased gradually up to 15^th^day (maximum33.5 fold) afterwards decreased in 25^th^and 35^th^days analysis. However, in Fe-sufficient condition, the expressionof *TaIds3* was detectable but almost constant at all stages (Fig. 5). The relative expression analysis reflected that Fe deficiency directly regulates the induction of *TaIds3* expression in Chinese Spring roots. The expression of *TaIds3* in Fe-deficient roots was much greater than Fe-sufficient roots at all stages (Fig. 5).

**Fig. 5.**
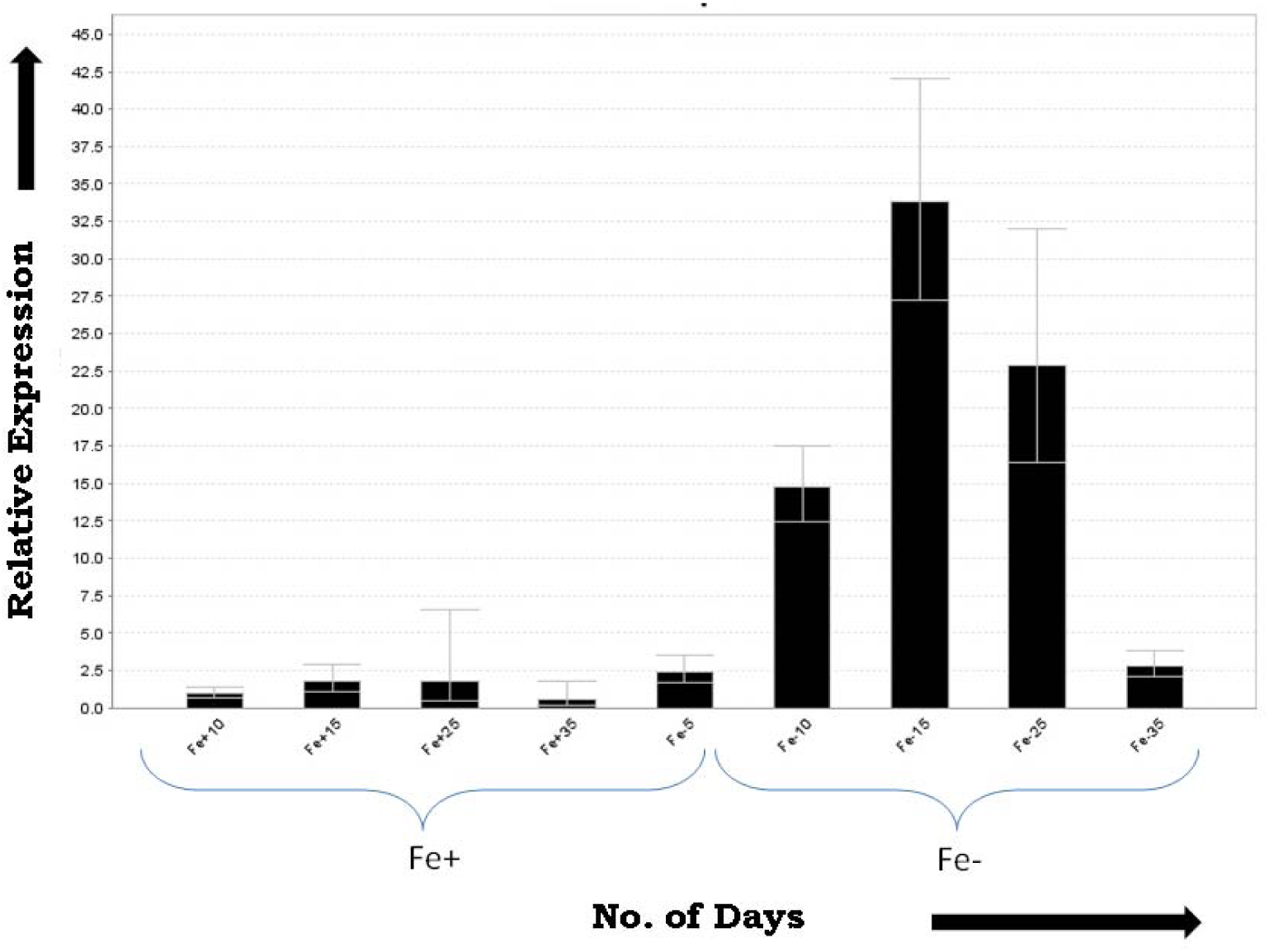
Relative expression level (change in fold) of *TaIds3* gene in roots of *T. aestivum* cv. Chinese Spring. Under Fe-sufficient and -deficient conditions at 5, 10, 15, 25 and 35 days after transplantation of seedlings in hydroponic condition. The means of three independent experiments are shown. Error bars indicate Standard Deviation.

### Molecular dynamics simulations

The molecular dynamics simulation concerned with stability and fluctuations in protein model of ids3 by analyzing 20nsec simulation trajectories. RMSD plot in Fig. 6B is show protein structure has RMSD between 1.3Å – 2.7Å. From the starting point (1nsec) of simulation, RMSD shows higher value between 3.4Å – 3.7Å, because protein structure has large loop region residues 4–46 that’s why it shows conformational changes between 1 to 3nsec. After 3nsec structure conformations are shows stable RMSD between 2.2Å – 2.7Å (Highlighted in red line) in 4 to 13nsec simulation time. After 13nsec RMSD shows some conformations changes due to second loop region 173–179 in protein and RMSD decrease 2.7Å to 1.6Å for 2 nsec (13–14nsec). After 15nsec RMSD remain constant and stable between 1.4Å – 1.6Åto till end of the simulation time (Highlighted in red line).

**Fig. 6.**
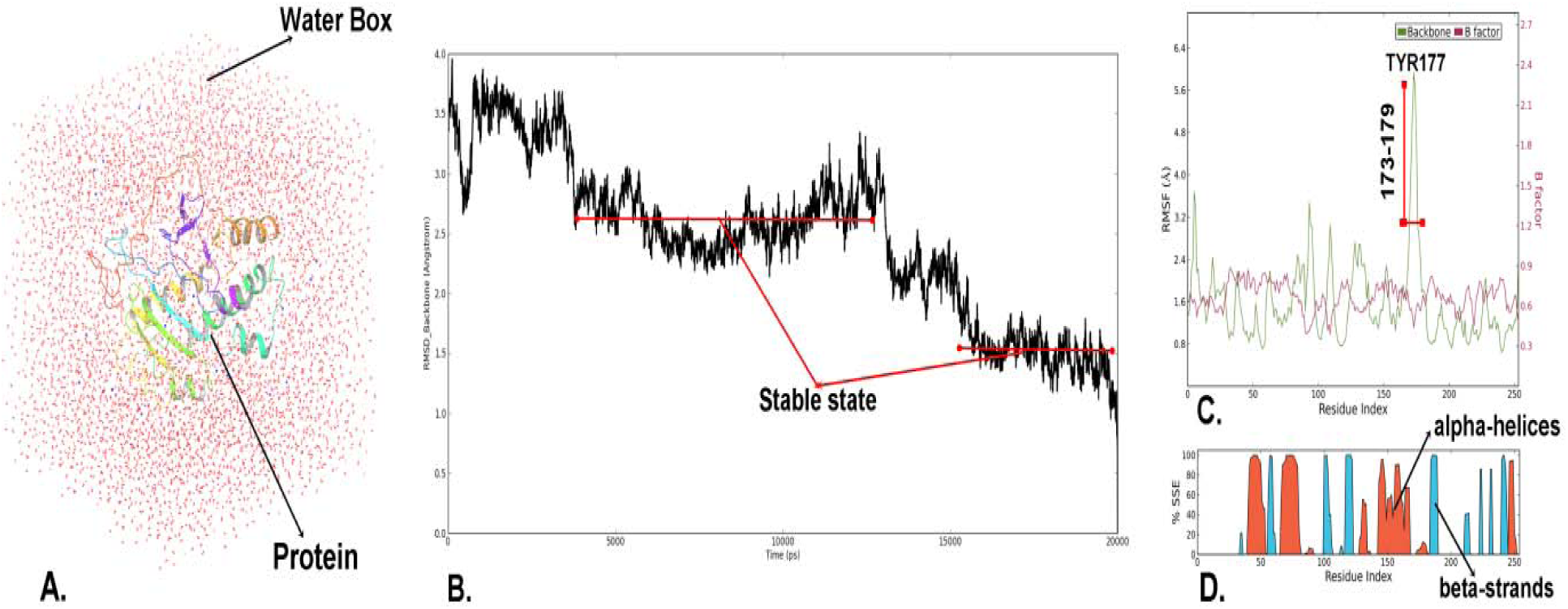
Molecular dynamics simulation results of HvIDS3 protein model. **A**. Simulation box with TIP3P water model, **B**. RMSD plot of IDS3 backbone atoms, **C**. RMSF plot with beta factor (experimental x-ray B-factor) of IDS3 backbone atoms and **D**. secondary structure elements of IDS3 protein model during simulation.

Fluctuations in IDS3 protein residues were analyzed by backbone atoms motions and local changes in secondary structure elements. RMSF plot Fig. 6C shows acceptable fluctuations under 2.4Å with compare to beta factor (0.5Å – 0.8Å) aspect one large fluctuations higher than 3Å between 173 – 179 residues range. This range comes in loop region of protein that’s why this region fluctuated during 13–14nsec simulation. Total 31.57% protein contributed in secondary structure (alpha helices=19.52% and beta strands =12.05%). SSE elements confirms that protein structure have stable conformations state aspect small loop (residues 173–179) that shows conformational changes during simulation. Based on acceptable RMSD around 1.5Å – 2.7Å and fluctuations between 1.0Å – 2.4Å confirms that protein has stable state during whole simulation time.

## Discussion

In the present study, we reported that*Ids3* gene, which encodes a dioxygenase that converts DMA to MA or MA to epiHMA, is present in hexaploid wheat. These results are contrary to a number of earlier reports showing absence of the *Ids3* gene(s) or presence of a mere pseudo gene in hexaploid wheat and its wild relatives (Nakanishi et al. 2000; Mori et al. 1990; Ma and Nomoto 1994) The bioinformatics analysis indicated that the functional wheat ortholog of *Ids3* gene might be present at chromosome 7AS, which is supported by the results obtained from physical mapping of the wheat ortholog in the telomeric region of chromosome 7AS. In the constructed phylogenetic tree, the close evolutionary relationship of *TaIds3_7AS* with *HvIds3* also supports the above results (Fig. 1). Our results are also supported by the location of most of the QTLs of high grain iron and zinc contents on chromosome 7A of different *Triticum* species (Tiwari et al. 2009; Peleg et al. 2009).The *Ids3* gene has been mapped on long arm of chromosome 4H of barley and long arm of chromosome 5R of rye using wheat-barley addition lines and wheat-rye addition lines respectively (Ma and Nomoto 1994; Mori and Nishizawa 1989; Ma et al. 1999).

The results of real-time RT-PCR showed the expression of wheat ortholog of *Ids3* gene in hexaploid wheat, which is strongly up-regulated under conditions of Fe-deficiency. According to the previous studies, during15-days of Fe starvation, the expression of *Ids3* was induced after three days of Fe-starvation in barley, which was gradually increased to a maximum level on the seventh day while the expression of *Ids3* gene was not observed in hexaploid wheat (Nakanishi et al. 2000). However, our results showed that the detectable expression of *Ids3* was observed five days after Fe-starvation and increased up to 15 days and thereafter its expression declined. So, the *Ids3* expression seemed to be more sensitive to Fe-deficiency in barley than in wheat. Similar expression pattern was reported for *Ids2* gene in barley. The *Ids2* expression in barley was hardly detectable at least for the first 7-days after the start of Fe-starved treatment (Okumura et al. 1994), while the expression of its ortholog in tobacco was clearly detectable within 3 days after Fe-starved treatment (Kobayashi et al., 2007). The induction of *Ids3* in barley is dependent on the availability of Fe and it is negatively correlated with the induction of NAS, NAAT and IDS3 enzymes and subsequently with secretion of MAs (Kobayashi and Nishizawa 2012; Higuchi et al. 1996; Kanazawa et al. 1998).These results strongly support the results of present investigation that the deficiency of Fe directly regulates the induction of *Ids3* expression in the roots of hexaploid wheat. Nakanishi et al. (1993) reported that the *Ids3* gene and its product detected only in Fe-deficient barley roots. Tolay et al. (2001) reported that as compared with diploid wheat (AA) and tetraploid wheat (AABB) species release higher amount of phytosiderophores under Fe deficiency. However, in case of Zn deficiency, diploid wheat species release higher amount of phytosiderophores than tetraploid wheat species. But in both the cases of deficiency, hexaploid wheat was found to secrete highest amount of phytosiderophores than diploid and tetraploid wheat species.This might be because the genes responsible for synthesis of phytosiderophores are expressed predominantly and effectively when three genomes are present together.On the basis of “Draft genome of the wheat A-genome progenitor *“Triticum urartu*” (Ling et al. 2013), homology analysis of the *Hordeum vulgare* IDS3 protein as a query with BLASTp exhibited 65% identity with *Triticum urartu* (Accession no. EMS 480341) and predicts the molecular function exactly similar as HvIDS3 protein i.e. dioxygenase and oxidoreductase activity (http://www.uniprot.org/uniprot/M7ZIY2). These results suggest the presence of IDS3 protein ortholog also in diploid wheat, which might be responsible for higher secretion of phytosiderophores in Fe and Zn deficient conditions. The good correlation between the protein and RNA data demonstrates that regulation of gene expression by concentration of Fe is at the level of transcription or RNA stability and not at the level of translation, as in the case of Fe-regulated genes in other systems. In spite of the presence of *Ids3* gene and its mRNA transcripts in hexaploid wheat genome, the less RNA stability might be the reason to produce the optimum amount of IDS3 peptide to be detectable in the root exudates.

In the present study, the *HvIds3* gene sequence was aligned against wheat survey sequence. BLASTn results showed 74% sequence similarity with IWGSC_chr7AS_ab_k71_contigs of chromosome 7AS in wheat genome having same conserved domain. Results indicate the presence of putative ortholog of *HvIds3* in the short arm of chromosome 7A of hexaploid wheat genome. Further, results of structural alignment of proteins between *HvIds3* and TaIds3ortholog by FATCAT server showing 57.25% similarity of *HvIds3* with putative wheat ortholog of *Ids3* indicated significant structural similarity in the proteins of both the genes which is an agreement with the observations of (Domingues et al. 2000). According to Domingues et al. (2000), if a protein has more than 40% sequence identity to another protein whose biochemical function is known and if the functionally important residue (for example, those in the active site of an enzyme) are conserved between the two protein, a reasonable working assumption can be made that the two proteins have a similar biochemical function. ProFunc analysis also identified the functional motifs of TaIDS3 and showed the close relationships of TaIDS3 to functionally characterized HvIDS3 protein. Even the subcellular locations of the gene in both wheat and barley was found in cytoplasm also suggests the presence of putative barley ortholog in wheat. Crystal structure of cereals IDS3 protein is not available in PDB yet. Moreover, high-throughput protein structure methods, such as yeast X-ray crystallography and NMR, cannot be easily applied to due to their high cost. Therefore, homology modeling for predicting RWP-RKs protein structure and function could provide alternative solutions. Homology modeling referred to as comparative or knowledge based modeling method, rely on detectable significant similarity between the query amino acid sequences and protein of know 3D structure (resolved by X-ray crystallography and NMR) and can be used to modelled the 3D structure of all the members of a protein family using a single representative 3D structure as an initiating point (Kumar et al. 2013; Kumar et al. 2016).The TaIDS3 sequence alignment with template is needed because Swiss-Model algorithm can discover structural information based on sequence. This alignment information is applied to simulation of target structure. Sequence alignment quality of target and template is important to predict a good homology model. Swiss-Model algorithm was predicted the 3D structure of TaIDS3 with good stero-chemical properties.

After protein structure modeling of ids3, we go for stability and local changes in structure analysis through molecular dynamics (MD) simulation for small time scale. Ids3 predicted model was place into cytoplasmic condition under TIP3P water molecules (Fig 6A). MD results suggest that predicted Ids3 protein model have stable state during simulation between 1.4Å – 2.7Åaverage rmsd (Fig. 6B). Root mean square fluctuations value between 1.4Å −2.4Å shows stable fluctuations in protein that compatible with experimental beta factor (Fig 5C). One large fluctuation in TYR177 residues that comes in loop region is reported between 3.2Å-5.7ÅRMSF. This large fluctuation doesn’t affect the local changes in secondary structure conformations. The percentage of secondary structure elements (31%) also shows that predicted model have stable conformation during simulation (Fig 6D).

## Conclusions

The present study first time reported the *HvIds3* ortholog in the hexaploid wheat genome and physically mapped on telomeric region of chromosome 7AS. The expression analysis reflected that Fe deficiency directly regulates the induction of *Ids3* gene in Chinese Spring roots. Modeling and MD simulations results of HvIDS3 revealed that identified IDS3 is stable at cell during Fee deficiency. Identified *TaIDS3* being utilizing by us in ongoing marker assisted program (MAS) to develop the Fe efficient wheat cultivar and boost up the biofortification program. However, extensive characterization and functional validation of *IDS3* gene in other cereals is also necessary to further explore their biological roles using different reverse genetics approaches like RNAi and VIGS.

## Author’s Contribution

SK and HSD designed and supervised the study. PM, UK and AK designed experiment and analyzed the data as a whole and wrote the manuscript. SM and PM collected samples for the analysis. AK and SK performed the molecular dynamics analysis. All authors read and approved the final manuscript.

## Acknowledgement

Thanks to Dr. B. S. Gill for providing the cytogenetic stocks for mapping.

